# Tuning the transglycosylation reaction of a GH11 xylanase by a delicate enhancement of its thumb flexibility

**DOI:** 10.1101/2020.12.19.423585

**Authors:** Kim Marneth, Hans van den Elst, Anneloes Cramer-Blok, Jeroen Codee, Hermen S. Overkleeft, Johannes M. F. G. Aerts, Marcellus Ubbink, Fredj Ben Bdira

## Abstract

Glycoside hydrolases (GH) are attractive tools for multiple biotechnological applications. In conjunction with their hydrolytic function, GH can perform transglycosylation reaction under specific conditions. In nature, oligosaccharides synthesis is performed by glycosyltransferase (GT). However, the industrial utilization of GT is limited by their instability in solution. A key difference between GT and GH is the flexibility of their binding sites architecture. In this report, we used the xylanase from *Bacillus circulans* (BCX) to study the interplay between active site flexibility and the transglycosylation reaction. Residues of the BCX thumb were substituted to increase the flexibility of the enzyme binding site. Replacement of the highly conserved residue P116 with glycine shifted the balance of the BCX enzymatic reaction toward transglycosylation. The effects of this point mutation on the structure and dynamics of BCX were investigated by NMR spectroscopy. The P116G mutation induces subtle changes in the configuration of the thumb and enhances the millisecond dynamics of the active site. Based on our findings, we propose the remodeling of the GH enzymes glycon site flexibility as a strategy to improve the transglycosylation efficiency of these biotechnologically important catalysts.

## Introduction

Oligosaccharides and their conjugates have found a wide range of pharmaceutical and biotechnological applications. For instance, in the food industry, oligosaccharides are used as prebiotics to improve the nutrition value of food by stimulating the growth of beneficial intestinal microflora.^[1]^ Oligosaccharide derivatives have multiple therapeutic and cosmetic uses.^[2]^ To fulfil the market demand, significant efforts were dedicated to the development of synthesis strategies of oligosaccharides with high yield and low-cost. Synthesis by chemical methods is tedious because it requires multiple protection and deprotection steps and gives low yields.^[3]^ A greener approach is to use enzymes to obtain a high yield of oligosaccharides with defined stereochemistry. Enzymatic synthesis can be achieved by glycosyltransferase (GT); however, these enzymes are sparse and unstable in solution, which makes them unsuitable for industrial use.^[4]^ An alternative is to use the ubiquitous family of glycoside hydrolases (GH).^[5]^

Retaining GH hydrolyze the glycosidic linkage using the “Koshland” double displacement mechanism with a catalytic nucleophile and acid/base for catalysis.^[6]^ The glycosyl-enzyme adduct is a common intermediate in hydrolysis and transglycosylation reactions. During the deglycosylation step, a water molecule or an alcohol, respectively, act as nucleophilic acceptors in the aglycone site of the enzyme, after being activated by the acid/base residue of the catalytic dyad (**Fig. 1A**).^[7]^ Therefore, the GH enzymes can perform carbohydrate synthesis and the balance between the two reactions is influenced by pH and temperature, as well as the availability of suitable acceptors like methanol and ethanol.^[8]^ However, the simultaneous presence of both activities represents a downside of this synthesis strategy, because newly formed products can get hydrolyzed in subsequent reactions. This bottleneck of the enzymatic carbohydrate synthesis was surmounted by the engineering of the GH enzymes to reduce or eliminate their hydrolytic activity. The new class of enzymes were termed glycosylsynthases (GS).^[5]^ Multiple GH engineering strategies were adopted, including the substitution of the catalytic residues and the use of fluorinated substrates.^[8]^ Introduction of mutations in the aglycon site has improved the binding affinity of acceptors molecules and thus the transglycosylation efficiency of the GH enzymes.^[9]^ Although GH and GT enzymes share similar active site architectures and topologies, they differ in flexibility. GT enzymes tend to be flexible, which is thought to be a requirement for the synthetic reaction.^[7b, 10]^ On the contrary, the rigidity of the GH appears to be necessary for their hydrolytic function.^[11]^ In an attempt to gain new insight into the role of flexibility in transglycosylation, we used xylanase from *Bacillus circulans* (BCX). BCX is a globular GH11 member exhibiting a conserved β-jelly-roll fold.^[12]^ The enzyme is remarkably rigid due to an extensive hydrogen bond network, restricting dynamics on the ps-ms NMR time scale.^[11b, 13]^ Recently, we found evidence that the enzyme induces a distortion of the substrate in the Michaelis complex that could facilitate the formation of the glycosyl-enzyme adduct.^[11b]^ The antiparallel ß-sheets of the β-jelly-roll of BCX shape a binding cleft that is 25 Å long, 9 Å deep and 4 Å wide. The cleft includes three (-) subsites (glycon binding site) and three (+) subsites (aglycon binding site, **Fig. 1B**).^[12]^ The glycosidic bond that is broken is located between the −1/+1 subsites with the substrate occupying at least the −2/−1 and +1 subsites, in accordance with the endo-catalytic mechanism. The overall shape of BCX fold is often compared to a right-hand fist, which includes a hand palm, thumb and fingers region. The PSIXG sequence motif of the thumb tip is highly conserved among the GH11 members, suggesting that this hairpin loop is important for the function.^[14]^ Molecular dynamics studies proposed that the modulation of the thumb flexibility might influence the substrate binding and product release from the enzyme binding cleft.^[15]^ Thus, to remodel the flexibility without interfering with the residues of the catalytic site, we generated BCX variants in which residues of the thumb were substituted with glycine or alanine. The effects of these point mutations on structure, dynamics and activity were probed. Paramagnetic NMR and relaxation dispersion (RD) NMR spectroscopic experiments showed that the mutation of the highly conserved residue P116 to a glycine induces subtle changes in the structure and dynamics of the thumb. The possible mechanism of the shift in activity toward transglycosylation as a consequence of the mutation is discussed.

**Figure 1:**
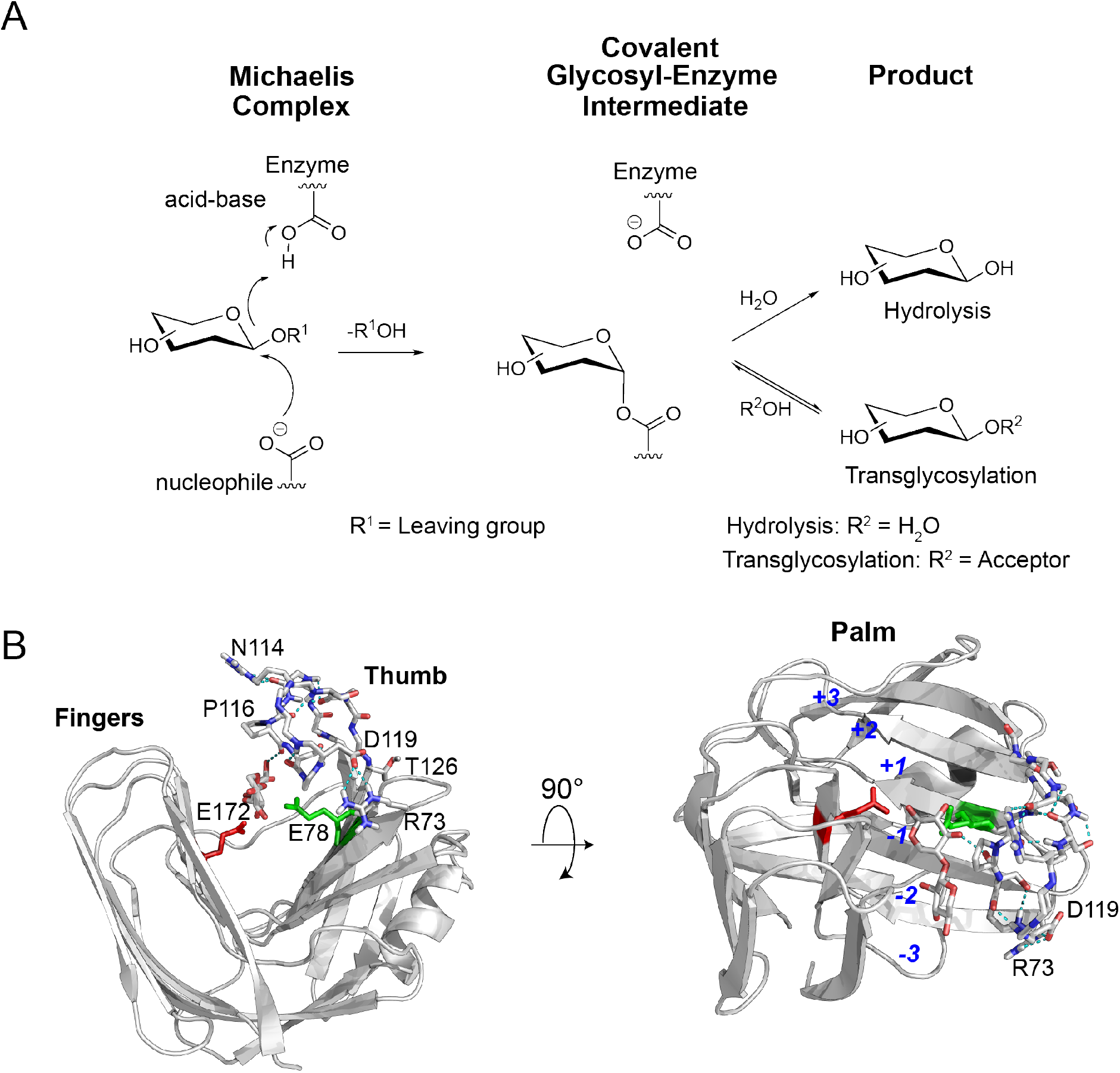
Catalytic mechanism of the GH enzymes and the fold topology of BCX. (A) The hydrolysis and the transglycosylation reactions of the GH enzymes. (B) ß-Jelly roll fold of BCX (PDB:1bcx)^[12]^ shown in grey cartoon. Residues of the thumb and the catalytic dyad (E78, E172) are shown in sticks. The internal hydrogen bonds of the thumb are shown in dashed cyan lines. The 2-fluoro-ß-xylobioside at the −1/−2 subsites is shown in sticks. Note the hydrogen bond between the O3 of the sugar and the backbone carbonyl of residue P116. In the right panel, the (+) aglycon subsites and the (-) glycon subsites are numbered in the enzyme binding cleft. Residues that form a salt bridge between the tip of the thumb and the palm are labelled.

## Results and discussion

### Design of the BCX mutants

A unique feature of the GH 11 family is the presence of a thumb loop that is key for the enzyme substrate turnover.^[14a]^ Residues of this loop are in contact with the substrate at the −1 and −2 subsites of the enzyme binding cleft (**Fig. 1B**).^[12]^ The thumb region displays a well-structured classical hairpin, containing a type I β-turn (residues 111-126 in BCX) with six internal hydrogen bonds (**Fig. 1B**). Multiple molecular dynamics simulations on different members of GH11 xylanase suggested the flexibility of this region and a putative role in substrate binding and product dissociation.^[15b, 16]^ However, in BCX, this region appears to be rigid with restricted dynamics on a wide range of NMR time scales.^[11b, 13]^ This difference in the thumb flexibility between GH11 members appears to be due to the difference in length and sequence of the thumb, although the PSIXG sequence motif of the thumb tip is highly conserved.^[14b]^ Thus, we aimed to introduce mutations on different positions of the thumb to enhance the flexibility of BCX binding cleft at the glycon site and probe their effects on the enzyme-catalyzed reactions. The first mutant was designed by substituting the residue P116 with a glycine. The almost complete conservation of this residue between all members of GH11 indicates its importance for the specific configuration of the thumb and the narrowing of the enzyme binding cleft for better selectivity towards non-decorated substrates.^[14a]^ The backbone carbonyl of P116 accepts a hydrogen bond from the O3 of the sugar substrate at the −1 subsite (**Fig. 1B**). By replacing P116 with a glycine it is possible that the rigidity within the thumb loop will be diminished, increasing flexibility. The hydrogen-bonding pattern of the thumb is interrupted by N114 leading to a sharp bend in the thumb structure (**Fig. 1B**). Additionally, it has been suggested that this residue represents a hinge point of the thumb in the GH11 family member, BsXynA.^[16]^ The configuration of the thumb is stabilized by a salt bridge between R73 of the palm and D119 of the thumb tip (**Fig. 1B**). Thus, to trigger conformational freedom of the thumb, we also substituted N114 and R73 with glycine and alanine, respectively. In BsXynA another mutation, D11F, allowed the capture in the crystalline state of the thumb in an open conformation with a relocation of its tip by more than 15 Å.^[17]^ Thus, we reasoned that the substitution of D11 with a glycine might also promote flexibility by excluding the bulky side chain of the amino acid from the BCX aglycon site. Finally, T126 is a potential hinge point that links the thumb to the palm.^[16]^ Substitution of T126 with proline in BsXynA improved the hydrolysis activity, possibly by increasing the rigidity of the thumb.^[17]^ Thus, as a control variant, we substituted T126 with a proline in BCX.

### Effects of the point mutations

To test whether the introduced mutations in BCX influenced the thermal stability, the melting temperature (Tm) was measured with a thermofluor assay.^[18]^ BCX WT has a Tm of 58 °C. The mutations reduced the Tm by 1 °C to 7 °C (**Fig. 2A**). The most significant destabilization effect was observed for mutations P116G and R73A, possibly due an increase in local flexibility. The melting temperature of mutant T126P is similar to the wild type, suggesting a minor effect on the protein rigidity. All the mutants have a Tm above 50 °C, so their hydrolytic activity was measured at 40 °C, the optimum temperature for WT BCX, using an end-point enzymatic assay with the artificial substrate 4-methylumbelliferone-xylobioside (4MU-X2). Most mutants show a reduction of hydrolytic activity of 20 to 30%. However, for BCX P116G it is reduced by 90% (**Fig. 2A**). These findings agree with previous reports in which the replacement of residues in BsXynA thumb by either glycine or proline, to modulate the flexibility, reduced the activity of the enzyme.^[17]^ Nevertheless, the activity enhancement observed for BsXynA T126P is not observed in BCX.

**Figure 2:**
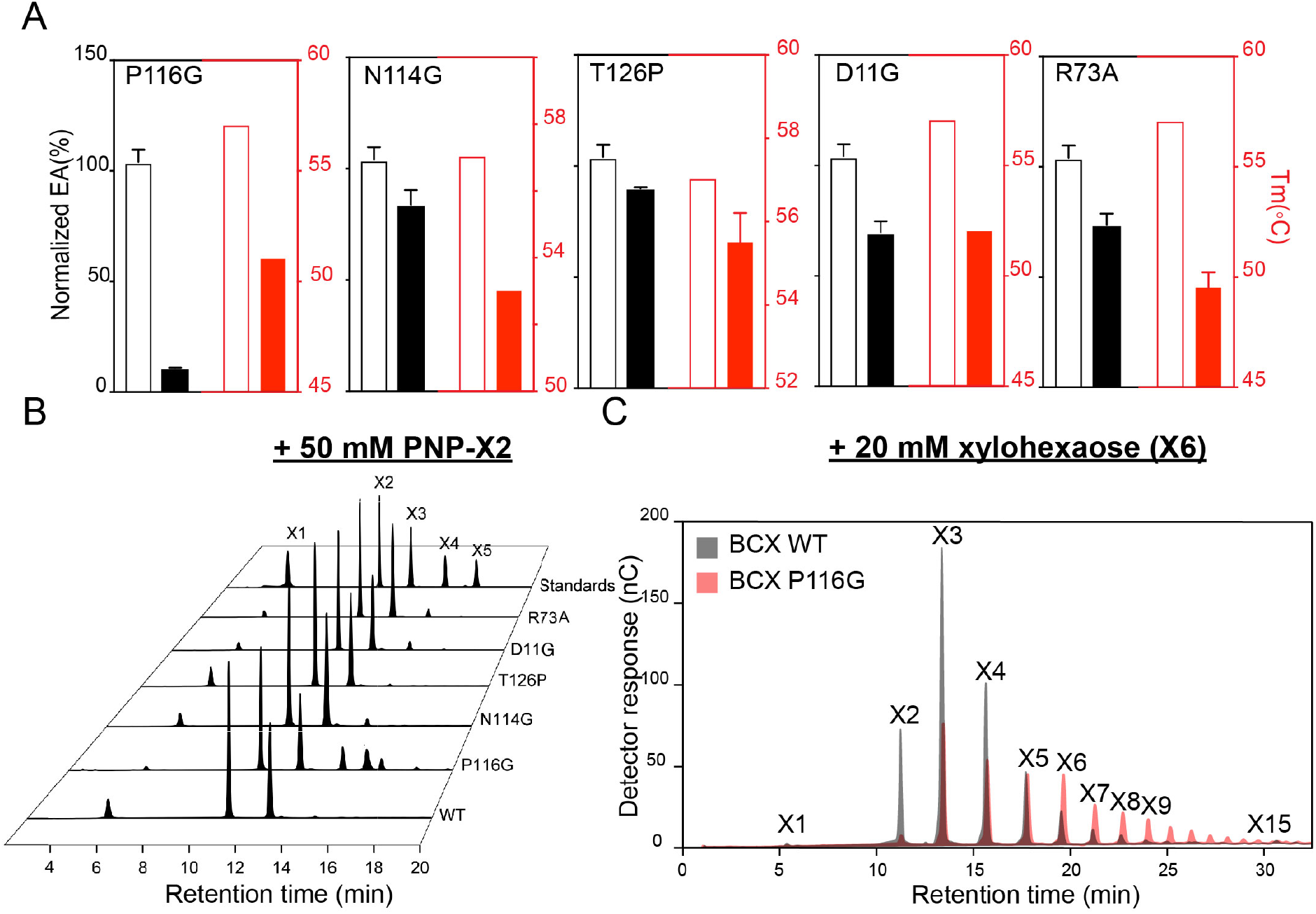
Effects of the thumb point mutations on the BCX activity and stability. (A) In the black and red panels, the 4MU-X2 hydrolytic enzymatic activity (EA) and melting temperatures, respectively, are depicted in percent of BCX WT (open bars) vs. the mutants (filled bars). (B) Ion exchange chromatograms of the transglycosylation products of the BCX variants. The reaction was conducted in the presence of 50 mM PNPX2 at 30% DMSO. A mixture of xylosides with different DP was used as standard. (C) An overlay between the ion exchange chromatograms of the BCX WT (in grey) and the BCX P116G (in red) transglycosylation products. The reaction was performed in the presence of 20 mM X6 in water (without DMSO) incubated for 30 min at 30 °C.

Although the trans-glycosylation activity has been reported for many GH families under specific conditions, it remains unknown if it also occurs for GH11 members in the presence of oligosaccharides acceptors. Previously, transglycosylation was not observed in BCX at high concentrations of its natural and artificial substrates.^[19]^ However, this activity was reported for the GH11 xylanase from *Thermobacillus xylanilyticus* in the presence of polyphenolic acceptors.^[9d]^ To test whether BCX is capable of transglycosylation in the presence of oligosaccharides acceptors, the enzyme was incubated with 50 mM of p-nitrophenyl-β-D-xylobioside (PNP-X2) at 30% DMSO for 30 min at 30°C. The oligosaccharides of the reaction mixture were analyzed by TLC (**Fig. S1A)**, showing several bands that suggest the formation of products with different degree of polymerization (DP). Separation by ion exchange HPLC revealed that the reaction products comprised the X1, X2 and X3 oligosaccharides (**Fig. 2B**). Although X2 might result from the hydrolysis of the glycosidic bond in PNP-X2, X3 and X1 can only occur from a transglycosylation reaction and/or the hydrolysis of newly formed products with DP ≥ 3. The observed transglycosylation reaction of BCX might be due to the presence of DMSO, which is known to reduce the activity of water molecules, leading to the accumulation of the enzyme-glycosyl intermediate.^[8a]^ This promotes the transfer of oligosaccharides units from the glycon site to the acceptor in the aglycon site of the enzyme. Therefore, we have tested the transglycosylation activity of BCX with xylohexaose (X6) at 20 mM concentration in water (without DMSO). HPLC analysis identified oligosaccharides with DP7, DP8 and DP9 in the reaction mixture (**Fig. 2C**). Also, shorter oligosaccharides (DP1, DP2 and DP3) are present due to the simultaneous hydrolysis activity of the enzyme. After establishing the presence of transglycosylation activity in BCX WT, we have tested the effect of the thumb mutations on this activity. Similar to their effects on the hydrolysis reaction, most of the thumb mutants did not have a major impact on the transglycosylation reaction of BCX (**Fig. 2B**). However, for mutant P116G, an accumulation of longer oligosaccharides was observed, ranging from three to four xylose units when PNP-X2 is used as a substrate (**Fig. 2B**). Additionally, the amount of X1 and X2 in the reaction mixture, which are products of hydrolysis, is reduced. The HPLC analysis of the reaction mixture of BCX P116G in the presence of 20 mM X6 revealed a significant shift in the enzyme activity towards the synthesis of longer oligosaccharides (up to DP15), compared to the wild type enzyme (**Fig. 2C**). Clearly, the hydrolytic activity of this variant is reduced.

### Effect of the P116G mutation on the protein structure and dynamics

The observed shift in activity specificity of BCX by the P116G substitution might be due to changes in the structure and dynamics of the enzyme. To investigate the effect of the point mutation on BCX structure, we recorded the ^1^H-^15^N HSQC spectra of the wild type and the mutant at 30°C and calculated the chemical shift perturbations (CSP). The P116G mutation results in small CSP localized in and around the thumb region with minor effects on residues of the fingers (**Fig. 3A**). These CSP report on changes in the chemical environment of the backbone amides, which could be due to the proline substitution and/or subtle structural rearrangements. Pseudocontact shifts (PCS) are sensitive probes to structural changes within a protein. PCS of the BCX WT and the P116G mutant were obtained by introducing two cysteine residues (T109C/T111C)^[11b]^ for attachment of the paramagnetic tag CLANP-5-Yb^3+^ or the diamagnetic control tag CLANP-5-Lu^3+^.^[20]^ For both WT and mutant BCX more than 130 PCS were obtained, by comparison of ^15^N-^1^H HSQC spectra of paramagnetic and diamagnetic samples, which were used to determine the position, the size, and the orientation of the anisotropic component of magnetic susceptibility tensor (Δχ) tensor. The data for WT BCX were fitted to the crystal structure with PDB ID 2bvv^[21]^ and the lanthanoid position was ~ 8 Å from the double cysteines Ca, in line with the previously reported distances (**Fig. 3A**).^[20]^ The magnitudes of the axial and rhombic components of the tensor were determined to be 8.15 ± 0.03 × 10^-32^ m^3^ and 2.36 ± 0.04 × 10^-32^ m^3^, respectively. An excellent agreement between the experimental and the predicted PCS was obtained with a Qa value (eq. S2) of 0.024 (**Fig. S1B**). These results obtained at 30°C are in line with the reported ones at 20°C.^[11b]^ Molecular dynamics (MD) simulations on BCX at different temperatures have suggested a temperature-dependent movement of the thumb region.^[22]^ A full opening of thumb was also captured in the crystalline state for the BsXynA D11F mutant.^[17]^ The occurrence of an open – closed motion of the thumb has also been proposed by several MD simulation studies on other GH11 members.^[14b, 15b, 16]^ However, the PCS data do not support such motion of the thumb in BCX and rather suggest that the conformation of the protein is the same at 20°C and 30°C (**Fig. S1B**).^[11b]^ Similarly, it was found using PCS that in mimics of the structures of intermediates of the catalytic cycle no large changes in the backbone occurred either.^[11b]^ Thus, we suggest that a full opening in the thumb does not occur and is not required for the function of BCX.

**Figure 3:**
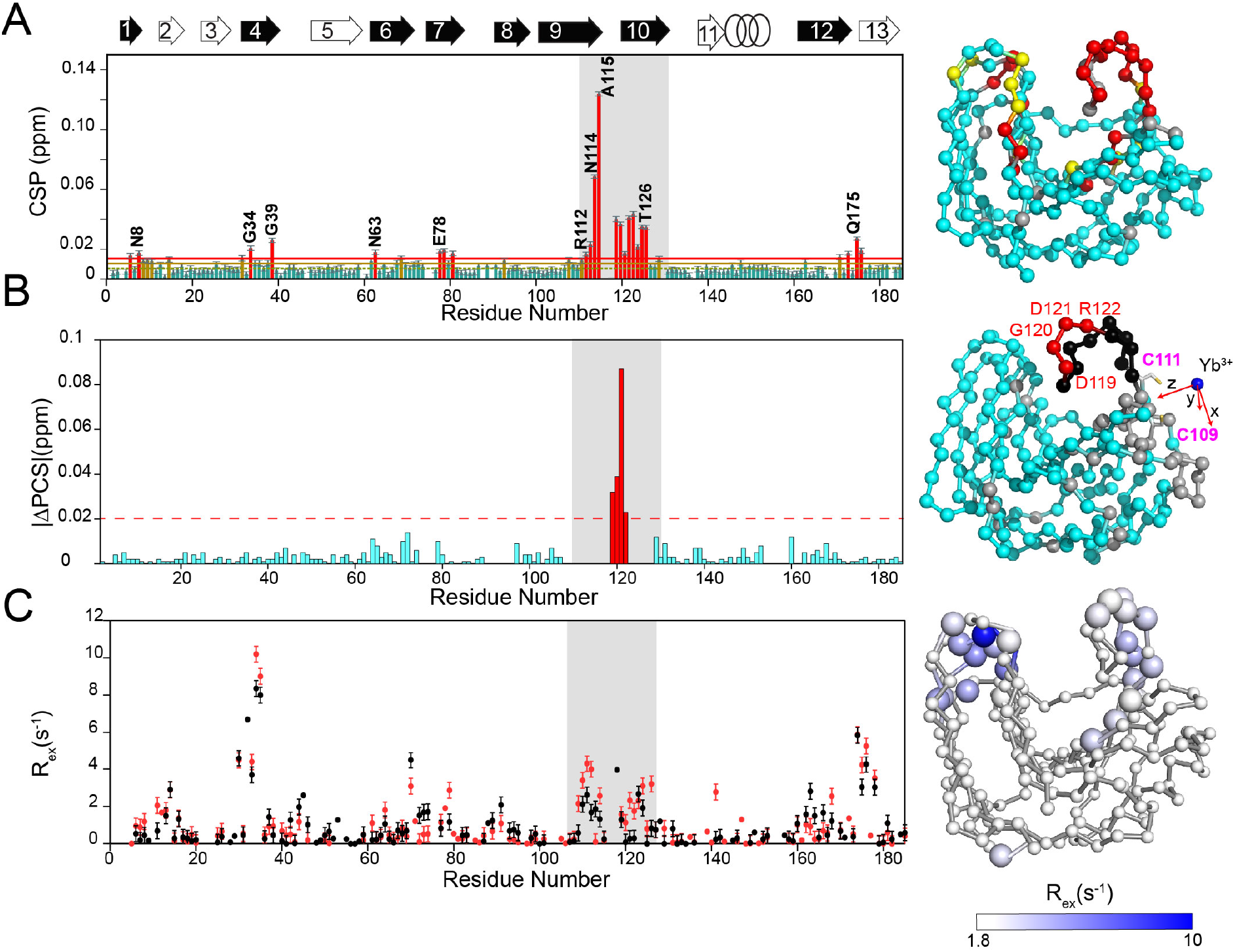
The effects of the P116G mutation on BCX structure and dynamics. (A) Analysis of the weighted average CSP between BCX P116G and WT. Resonances with CSP more than two (red line) or one (yellow line) standard deviation (SD) from the 10% trimmed mean (green dashed line) are labelled and shown in red, yellow, and green bars, respectively. In the right panel, amide nitrogens with a CSP > 1 and 2 SD are shown in spheres on the BCX structure (PDB:2bvv)^[21]^ and coloured in red and yellow, respectively. Nitrogens with no data are shown in grey. (B) The ļΔPCSsļ between BCX P116G WT are plotted versus residue number. ļΔPCSļ > 0.02 (red dashed line) are shown in red bars. In the right panel, amide nitrogens of the residues with ļΔPCSļ > 0.02 are mapped on the BCX structure in red spheres and labelled. Nitrogens with not data in the thumb region and elsewhere are depicted in black and grey, respectively. The blue sphere indicates the position of the lanthanide ion (Yb^3+^). The x, y, and z axes of the Δχ tensor are indicated in red arrows. The cysteine residues at which CLANP-5 was attached are shown in sticks and labelled. (C) The experimental *R*ex of the BCX WT (black dots) and P116G (red dots) versus residue number. In the right panel, the amide nitrogens of BCX P116G with *Rex* ≥ 1.8 s^-1^ are mapped on the BCX structure, shown in large spheres and coloured with a white/blue colour gradient. Nitrogens without significant ms dynamics are shown in small spheres. Above the top panel the BCX secondary structure is shown by black arrows for β-strands of sheet A and in white ones for sheet B and the α-helix in rings. The thumb region is highlighted with grey rectangles in the panels.

For BCX P116G, several PCS of the amide nuclei of the thumb could not be determined due to line broadening caused by paramagnetic relaxation enhancement (PRE).^[23]^ The PCS of the thumb residues D119, G120, D121 and R122 could be measured, however (**Fig. 3B)**. The broadening suggests proximity in the spatial configuration of the mutated thumb relative to the paramagnetic centre. The absolute values of the differences between the observed PCS of the BCX WT and P116G mutant (|ΔPCS|) indicate a highly localized structural change in the thumb of the protein (**Fig. 3B, S1C**). The rest of the backbone of the protein is unperturbed.

Structural motions in the millisecond time (ms) scale can influence substrate turnover by the enzyme in different ways. It can cause the exclusion of water molecules from the active site, optimize the position of either the substrate or the catalytic residues in the protein and assist the release of the product from the binding site.^[24]^ Therefore, to interrogate the effects of the P116G mutation on the ms dynamics of BCX, we performed a relaxation dispersion TROSY-CPMG experiment. Previously we have observed that BCX WT exhibits minor chemical exchange (Rex) for some residues remote from the catalytic site.^[11b]^ The P116G mutation slightly enhanced the ms chemical exchange of the enzyme binding cleft for residues of the thumb and fingers regions at the glycon site (**Fig. 3C**).

In summary, these data indicate that the P116G mutation induces a structural change and somewhat enhanced ms dynamics localized to the thumb region at the glycon site of BCX. A conformational change in the loop may expose the hydrophobic interface between the thumb and the underlying palm, which could explain the large decrease in Tm.

## Conclusion

GH enzymes are attractive tools in green chemistry for the synthesis of biotechnologically relevant oligosaccharides.^[1a, 2a]^ Several approaches were used to improve their synthesis efficiency by focusing on the optimization of the reaction conditions (pH, T°, organic solvent, acceptors) and the enhancement of the acceptor affinity toward the enzymes aglycon sites.^[5, 8a]^ Although these strategies proved to be successful in many GH, the role of enzymes binding site dynamics and flexibility in transglycosylation remained unexplored.

In this report, we attempted to shed light on the interplay between fold flexibility and the trans-reaction using a GH11 xylanase from *B. circulans* (BCX) as a benchmark due to its distinctive fold rigidity^[11b]^. We found that the substitution of the conserved residue P116 of the thumb with a glycine induced subtle changes in the ms dynamics and conformation at the glycon site. The mutant also exhibits a 10-fold reduction in hydrolysis activity, resulting in a shift in the balance between hydrolysis and transglycosylation activity toward the latter. It is possible that activation of the nucleophilic water molecule is less efficient, because of a slight displacement of the water molecule relative to the acid/base catalytic residue. The extensive network of ordered water molecules can transfer effects of the mutation throughout the active site. Alternatively, the glycosyl adduct, which is in direct contact with P116, might change position or conformation slightly, making the hydrolytic attack more difficult. For the alternative reaction, with a xylose acceptor, the rate reduction is less or absent, resulting in more transglycosylation and less hydrolysis. Nevertheless, the BCX P116G variant retains hydrolytic activity against its new synthesized products. Therefore, further investigations and protein design are required to eliminate this undesired reaction.

In conclusion, our study provides new insights into the possible role of fold flexibility in the transglycosylation reaction and propose the modulation of the rigid binding site characteristic of the GH enzymes as a strategy to engineer more efficient transglycosylation catalysts.

## References

[1a] J. A Linares-Pasten, A. Aronsson, E. Nordberg Karlsson, Curr. Protein. Pept. Sci. 2018, 19, 48–67;

[1b] J. Park, J. Yu, Y. Shin, H. Shin, S. Lee, K. Park, J. Microbiol. Biotechnol. 1992, 20, 237–242;

[1c] T. Oku, S. Nakamura, Pure Appl. Chem 2002, 74, 1253–1261.

[2a] W. von Rybinski, K. Hill, Angew. Chem. Int. Ed. Engl. 1998, 37, 1328–1345;

[2b] L. Greffe, L. Bessueille, V. Bulone, H. Brumer, Glycobiology 2005, 15, 437–445;

[2c] X. Vecino, J. Cruz, A. Moldes, L. Rodrigues, Crit. Rev. Biotechnol. 2017, 37, 911–923;

[2d] C. Duarte, E. J. Gudiña, C. F. Lima, L. R. Rodrigues, AMB express 2014, 4, 40;

[2e] E. J. Gudiña, E. C. Fernandes, J. A. Teixeira, L. R. Rodrigues, RSC Advances 2015, 5, 90960–90968.

[3] S. P. Douglas, D. M. Whitfield, J. J. Krepinsky, J. Am. Chem. Soc. 1991, 113, 5095–5097.

[4] S. Prapulla, V. Subhaprada, N. Karanth, Adv. Appl. Microbiol., volume 47, 2000.

[5] L. F. Mackenzie, Q. Wang, R. A. J. Warren, S. G. Withers, J. Am. Chem. Soc. 1998, 120, 5583–5584.

[6] D. Koshland Jr, Biol. rev. 1953, 28, 416–436.

[7a] S. Withers, Carbohydr. Polym. 2001, 44, 325–337;

[7b] V. Rojas-Cervellera, A. Ardèvol, M. Boero, A. Planas, C. Rovira, Chem-Eur. j. 2013, 19, 14018–14023.

[8a] N. H. Abdul Manas, R. Md. Illias, N. M. Mahadi, Crit. Rev. Biotechnol. 2018, 38, 272–293;

[8b] S. M. Hancock, M. D. Vaughan, S. G. Withers, Curr. Opin. Chem. Biol. 2006, 10, 509–519.

[9a] J. Durand, X. Biarnés, L. Watterlot, C. Bonzom, V. Borsenberger, A. Planas, S. Bozonnet, M. J. O’Donohue, R. Fauré, ACS Catalysis 2016, 6, 8264–8275;

[9b] P. Lundemo, E. N. Karlsson, P. Adlercreutz, Appl. Microbiol. Biotechnol. 2017, 101, 1121–1131;

[9c] N. N. Aronson, B. A. Halloran, M. F. Alexeyev, X. E. Zhou, Y. Wang, E. J. Meehan, L. Chen, Biosci. Biotechnol. Biochem. 2006, 70, 243–251;

[9d] C. Brusa, N. Belloy, D. Gérard, M. Muzard, M. Dauchez, R. Plantier-Royon, C. Rémond, J. Biotechnol. 2018, 272, 56–63.

[10] J. Nyirenda, S. Matsumoto, T. Saitoh, N. Maita, N. N. Noda, F. Inagaki, D. Kohda, Structure 2013, 21, 32–41.

[11a] A. White, D. Tull, K. Johns, S. G. Withers, D. R. Rose, Nat. Struct. Biol. 1996, 3, 149–154;

[11b] F. B. Bdira, C. A. Waudby, A. N. Volkov, S. P. Schröder, E. Ab, J. D. Codée, H. S. Overkleeft, J. M. Aerts, H. van Ingen, M. Ubbink, Angew. Chem. Int. Ed. Engl., 2020.

[12] W. W. Wakarchuk, R. L. Campbell, W. L. Sung, J. Davoodi, M. Yaguchi, Protein sci. 1994, 3, 467–475.

[13] G. P. Connelly, S. G. Withers, L. P. Mcintosh, Protein sci. 2000, 9, 512–524.

[14a] G. País, V. Tran, M. Takahashi, I. Boukari, M. J. O’Donohue, Protein Eng. Des. Sel. 2007, 20, 15–23;

[14b] G. País, J.-G. Berrin, J. Beaugrand, Biotechnol. Adv. 2012, 30, 564–592.

[15a] N. N. Mhlongo, M. Ebrahim, A. A. Skelton, H. G. Kruger, I. H. Williams, M. E. Soliman, RSC advances 2015, 5, 82381–82394;

[15b] G. País, J. Cortés, T. Siméon, M. J. O’Donohue, V. Tran, Comput. Struct. Biotechnol. J. 2012, 1, e201207001.

[16] D. S. Vieira, R. J. Ward, J. Mol. Model 2012, 18, 1473–1479.

[17] A. Pollet, E. Vandermarliere, J. Lammertyn, S. V. Strelkov, J. A. Delcour, C. M. Courtin, Proteins 2009, 77, 395–403.

[18] U. B. Ericsson, B. M. Hallberg, G. T. DeTitta, N. Dekker, P. Nordlund, Anal. Biochem. 2006, 357, 289–298.

[19] M. L. Ludwiczek, M. Heller, T. Kantner, L. P. McIntosh, J. Mol. Biol. 2007, 373, 337–354.

[20] P. H. Keizers, A. Saragliadis, Y. Hiruma, M. Overhand, M. Ubbink, J. Am. Chem. Soc. 2008, 130, 14802–14812.

[21] G. Sidhu, S. G. Withers, N. T. Nguyen, L. P. McIntosh, L. Ziser, G. D. Brayer, Biochemistry 1999, 38, 5346–5354.

[22] D. S. Vieira, L. Degrève, R. J. Ward, Biochim. Biophys. Acta. Gen. Subj 2009, 1790, 1301–1306.

[23] G. M. Clore, in Methods in enzymology, Vol. 564, Elsevier, 2015, pp. 485–497.

[24] K. Henzler-Wildman, D. Kern, Nature 2007, 450, 964–972.

