## Supplemental information for "Tuning the transglycosylation reaction of a GH11 xylanase by a delicate enhancement of its thumb flexibility"

### Materials and Methods

#### Materials

4-methylumbelliferone-xylobioside (4MU-X2), p-nitrophenyl-β-D-xylobioside (PNP-X2), xylobiose (X2) and xylohexaose (X6) were purchased from Megazyme.

#### Methods

**Mutagenesis**

BCX P116G, N114G, T126P, D11G, R73A, BCX T109C/T111C and BCX P116G T109/T111C mutants were prepared by site-directed mutagenesis using Quik-Change protocol.

**BCX variants overexpression and purification.**

BCX variants were overexpressed in E. coli and purified as previously described[1].

**BCX enzymatic activity:**

The relative hydrolytic activity of the enzyme was tested by an end-point assay, using the 4MU-X2 substrate. Typically, 5 μM of enzyme was incubated with 20 mM 4MU-X2 in 25 mM sodium acetate pH 5.7 for 1 hour at 40°C, in duplicate. The reaction was stopped by adding 2.5 mL of 0.3 M glycine buffer pH 10.6. The fluorescence emission of the 4MU product was measured at 490 nm using an excitation wavelength of 370 nm on a FP-800 series fluorometer SHIMADZU.

For the transglycosylation reaction, 5 μM of the enzyme was incubated with 50 mM PNP-X2, in 25 mM sodium acetate buffer, 30% of DMSO or with 20 mM X6 in 25 mM sodium acetate buffer at pH 5.7 for 30 min at 30°C. The reaction was stopped by flash freezing the mixture in liquid nitrogen. The transglycosylation products were separated on a TLC silica gel 60 F254. The mobile phase was an acetonitrile:H2O (1:9) solution. The products were visualized by spraying the TLC membrane with 20% sulfuric acid solution and heated with a hot air dryer.

**Ion exchange HPLC of the transglycosylation products:**

The transglycosylation products were analyzed by a Dionex ICS-5000 HPLC system (Thermo Scientific) with a ICS-5000 DC electrochemical detector. The reaction samples were mixed with 50 mM NaOH and 20 uL were injected into a Dionex CarboPac PA20 3x150mm column with a Dionex CarboPac PA20 3x30mm Guard. The oligosaccharides we eluted by applying a sodium acetate gradient from 0 to 50 mM for 20 min with a flowrate of 0.5 ml/min.

**NMR spectroscopy**

The $^{15}N$-$^{1}H$ HSQC spectra of BCX WT and P116G were recorded on a Bruker Avance III 600 spectrometer in 25 mM sodium acetate (pH=5.7), 6% $D_2O$ at 30°C and assigned with the published data[1] and curated with PCS (see below). NMR titrations of $^{15}N$ BCX WT and P116G (100 μM) with an increasing X2 concentration (100 μM-20 mM) was performed under the same conditions. The CSP were calculated according to equation (S1), using as references the spectra of the free proteins.

$$\mathrm{CSP} = \sqrt{\Delta\delta_H{}^2 + \frac{\Delta\delta_N{}^2}{25}} \quad \text{(S1)}$$

**Paramagnetic NMR spectroscopy**

Paramagnetic NMR samples were prepared as described previously. [1]. The $^{1}$H-$^{15}$N HSQC spectra of BCX WT and P116G were recorded on a Bruker Avance III 600 spectrometer at 30°C, processed by TopSpin 3.5 and analyzed with Ccpnmr analysis.[2] The chemical shift differences between the resonances in the spectra of paramagnetic and diamagnetic samples were taken as the PCS. The obtained PCS of the BCX WT were fitted to the protein crystal structure (PDB ID:2bvv) using Numbat,[3] and $\Delta\chi$ was determined. The quality of the fit was quantified by the $Q_a$ factor, calculated by the equation (S2)[4]:

$$Q_a = \sqrt{\frac{\sum_i (PCS_{obs} - PCS_{calc})^2}{\sum_i (|PCS_{obs}| + |PCS_{calc}|)^2}} \quad \text{(S2)}$$

Note that the nominator sums the experimental and calculated PCS, contrary to the standard Q factor.

**NMR relaxation dispersion and CPMG data analysis**

$^{15}$N RD-CPMG data set were recorded at 20°C at a field strength of 19.9 T on a Bruker Avance HD spectrometer equipped with a TCI-Z-GRAD triple resonance cryoprobe, as previously reported. [1] The constant time relaxation delay, $\tau_{CPMG}$, was set to 50 ms. Dispersion profiles comprised ~ 20 $\nu_{CPMG}$ frequencies, recorded in an interleaved manner, with values ranging from 20 to 1000 Hz. Errors were estimated based on repeat measurements at two $\nu_{CPMG}$ frequencies. The experimental $R_{ex}$ were calculated using the formula $R_{ex}$= $R_{2,eff\,(20\ Hz)}$- $R_{2,eff\,(1000Hz)}$.

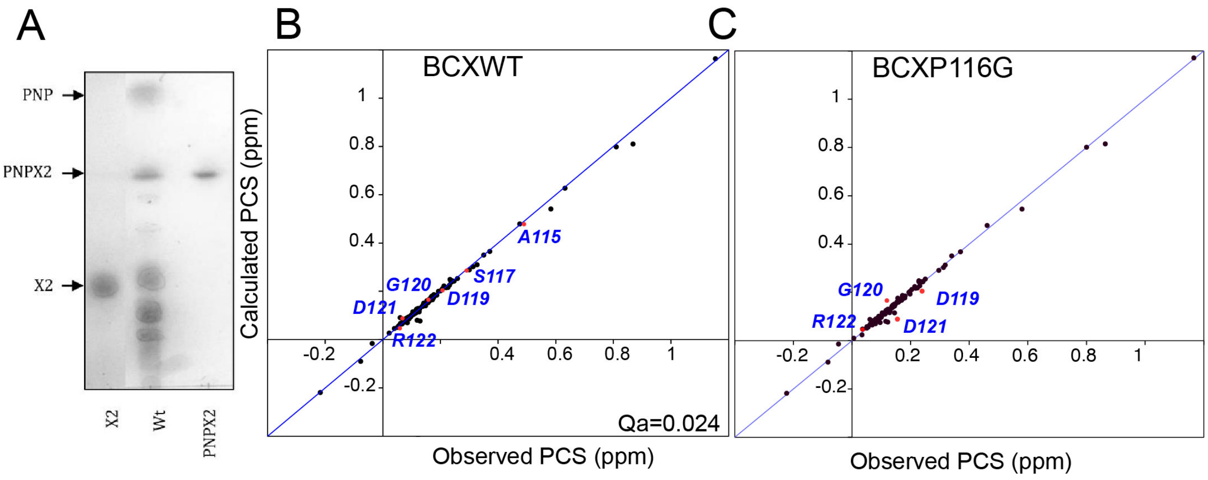


**Figure S1:** (A) TLC membrane of the transglycosylation products of the BCX WT in the presence of PNP-X2. (B) and (C) Correlation plot of the calculated vs. observed PCS of BCX WT and BCXP116G after fitting to the crystal structure BCX (PDB: 2bvv). Correlation points of the thumb residues are highlighted in red and labelled.


[1] F. B. Bdira, C. A. Waudby, A. N. Volkov, S. P. Schröder, E. Ab, J. D. Codée, H. S. Overkleeft, J. M. Aerts, H. van Ingen, M. Ubbink, Angew. Chem. Int. Ed. Engl., **2020**.

[2] W. F. Vranken, W. Boucher, T. J. Stevens, R. H. Fogh, A. Pajon, M. Llinas, E. L. Ulrich, J. L. Markley, J. Ionides, E. D. Laue, *Proteins* **2005**, *59*, 687-696.

[3] C. Schmitz, M. J. Stanton-Cook, X.-C. Su, G. Otting, T. Huber, *J. Biomol. NMR,* **2008**, *41*, 179.

[4] Q. Bashir, A. N. Volkov, G. M. Ullmann, M. Ubbink, *J. Am. Chem. Soc.* **2010**, *132*, 241-247.
